# Concomitant chemotherapy increases radiotherapy-mediated DNA-damage in peripheral blood lymphocytes

**DOI:** 10.1101/540575

**Authors:** Yvonne Lorat, Jochen Fleckenstein, Patric Görlinger, Christian Rübe, Claudia E. Rübe

## Abstract

53BP1-foci detection in peripheral blood lymphocytes (PBLs) by immunofluorescence microscopy (IFM) is a sensitive and quantifiable DNA-double-strand-break (DSB) marker. In addition, high-resolution transmission-electron-microscopy (TEM) with immunogold-labeling of 53BP1 and DSB-bound phosphorylated Ku70 (pKu70) can be used to determine the progression of the DNA-repair process. Here, we analyzed whether different modes of irradiation influence the formation of DSBs in the PBLs of patients with cancer, and whether accompanying chemotherapy influences the DSB-appearance.

We obtained 86 blood samples before and 0.1, 0.5 and 24 h after irradiation from patients with head and neck, or rectal cancers receiving radiotherapy (RT) or radio-chemotherapy (RCT). 53BP1-foci were quantified by IFM. In addition, TEM was used to quantify gold-labelled pKu70-dimers and 53BP1-clusters within euchromatin and heterochromatin of PBLs. During radiotherapy, persisting 53BP1-foci accumulated in PBLs with increasing numbers of administered RT-fractions. This 53BP1-foci accumulation was not influenced by irradiation technique applied (3D-conformal radiotherapy versus intensity-modulated radiotherapy), dose intensity per fraction, number of irradiation fields, or isodose volume. However, more 53BP1-foci were detected in PBLs of patients treated with accompanying chemotherapy. TEM analyses showed that DSBs, indicated by pKu70, were present for longer periods in PBLs of RCT-patients than in PBLs of RT-only-patients. Moreover, not every residual 53BP1-focus was equivalent to a remaining DSB, since pKu70 was not present at every damage site. Persistent 53BP1-clusters, visualized by TEM, without colocalizing pKu70 likely indicate chromatin alterations after repair completion, or possibly, defective repair. Therefore, IFM-53BP1-foci analyses alone are not adequate to determine individual repair capacity after irradiation of PBLs, as a DSB may be indicated by a 53BP1-focus but not every 53BP1-focus represents a DSB.

The level of DNA-damage during RT is influenced by the presence of accompanying chemotherapy.

## Introduction

The effects of radiotherapy (RT) in cancer treatment can be significantly enhanced by simultaneous chemotherapy [1]. The action of both seems to depend on their ability to induce mutagenic and clastogenic DNA damage, including crosslinks, strand-breaks, replication errors and base adducts [2, 3], which can induce cell death. Precise dose distributions to the planning target volume (PTV) by highly conformal techniques are critical for minimizing side effects to adjacent organs at risk.

DNA-damage-repair mechanisms protect against adverse effects of carcinogenic therapies. While the likelihood of RT-induced side effects in organs at risk can be reliably assessed by dosimetric calculations, peripheral blood lymphocytes (PBL), especially in patients who received radio-chemotherapy (RCT), are exposed to an erratic amount of events that may cause DNA damage. Double-strand-break (DSB) repair is crucial for PBL survival following RT or RCT-induced DNA damaging effects. During non-homologous end joining (NHEJ), the major mammalian DSB-repair-pathway, the Ku70-Ku80-heterodimer recognizes DSBs and maintains the broken DNA-ends in close proximity until the DSB is rejoined [4]. In addition, the phosphorylated histone variant of H2AX, γH2AX, recruits repair proteins such as 53 binding protein 1 (53BP1) to the chromatin surrounding the DSB. Specific primary and fluorescent secondary antibodies against γH2AX and 53BP1 localized in DSB-repair-foci [5, 6] may be used as markers to quantify DSB repair by immunofluorescence microscopy (IFM).

Assuming that each γH2AX-or 53BP1-focus corresponds to one DSB, the number of foci in the nucleus can be applied to measure DNA-damage caused by radiation exposure [7–13]. PBLs are suitable to assess the DNA-damage-response of patients as peripheral blood samples can be taken repeatedly and at defined time points after irradiation. In addition, the hematopoietic system is radiosensitive; lymphocytes and their subpopulations well characterized in terms of their phenotype and function [14–17], and can be reliably isolated from blood [18]. Moreover, PBLs are in the resting state (G0) of the cell cycle [19, 20], thereby resulting in prolonged presence of DNA-damage [21–23].

The limited resolution in IFM for γH2AX and 53BP1 visualization does not provide full-scale information regarding individual DSB repair points and radiation sensitivity. Additionally, individual repair proteins of the Ku70-Ku80-heterodimer cannot be detected as their fluorescence is not sufficient to differentiate them from the background signal. The detection of both 53BP1 and the DSB-bound phosphorylated Ku70 (pKu70) would signal incomplete NHEJ-repair-sites. Here it was sought to assess the impact of both RT and RCT on the genomic integrity of PBLs. High-resolution transmission-electron-microscopy (TEM) with gold-labeled pKu70 and 53BP1 [24, 25], in addition to IFM, was used to determine the suitability of this analysis in assessing individual PBL radiation sensitivity in patients with different tumor entities (head and neck, or rectal cancers), isodose volumes, irradiation techniques (IMRT or 3D-CRT), and treatment approaches (RT or RCT).

## Materials and methods

### Patients and treatment conditions

This study was conducted in accordance with the Helsinki declaration and with approval of the local ethics committee (Aerztekammer des Saarlandes). All patients signed written informed consent forms. Patients meeting the following inclusion criteria were enrolled between March 2011 and May 2012: Aged between 18 to 80 years; Karnofsky index >70%; completely resected head and neck squamous cell cancer (oral cavity, oropharynx, hypopharynx, or larynx) with postoperative RT indicated with or without chemotherapy; or diagnosis of rectal cancer with an indication for neoadjuvant or adjuvant pelvic radiotherapy (with or without chemotherapy). Patients with previous RT or chemotherapy and those with distant metastases were excluded.

All patients underwent standard computed tomography-based RT-planning with 3D-conformal target volume delineation. IMRT with a pre-defined PTV arrangement of seven coplanar beam angles with 70 beam-segments and standardized objectives based on the ICRU Report 83 [26] and constraints for normal tissues (brainstem, spinal cord, parotid glands, esophagus) based on QUANTEC-data [27] with individualized clinical assessments was mandatory for patients with head and neck cancer. A 60 Gy reference dose was prescribed to primary tumor sites and lymph node metastases in cervical regions and 50 Gy to non-involved cervical and supraclavicular lymph node regions. Single doses were 2.0 Gy, once daily, 5 days a week. Concomitant chemotherapy, if prescribed, included two cycles of cisplatin (20 mg/m^2^ intravenously over 0.5 h, D1-5; D29-33) and two cycles of 5-fluorouracil (600 mg/m2 intravenously over 24 h, D1-5; D29-33). Patients with rectal cancer were treated in a prone position on a belly board by means of three 3D-conformal coplanar portals (0°; 90°; 270°) with a total reference dose of 50.4 Gy (optional 5.4 Gy boost to the primary tumor after 45 Gy) and a single dose of 1.8 Gy (once daily, 5 fractions/week). Concurrent neoadjuvant chemotherapy consisted of two cycles of 5-fluorouracil (1000 mg/m^2^ intravenously over 24 hours, D1-5; D29-33). Adjuvant 5-fluorouracil was administered as a continuous infusion of 225 mg/m2 (D1-38). All patients were irradiated with a linear accelerator (ONCOR™ or ARTISTE™) from Siemens (Erlangen, Germany) using photons of 6 MV for IMRT of head and neck cancers or 18 MV for 3D-conformal RT (3D-CRT) of rectal cancers. The analysis of RT-related parameters included assessment of blood volume contained within the 50%-isodose line (derived from volumetric computation of delineated blood vessels >1 cm in diameter), body volumes surrounded by the 10 Gy, 20 Gy, 30 Gy and 45 Gy isodose lines (*V10_iso_ − V45_iso_*) and coverage of the PTV (*D80, D90*).

### Blood sampling

For IFM-analysis, blood samples were collected from a cubital vein in heparin-containing vials at 37°C and diluted 1:2 with pre-warmed Roswell Park Memorial Institute (RPMI) 1640 medium (Biochrom; Berlin, Germany) for immediate processing. All patient samples were obtained immediately before and 0.5 h after the first RT-fraction (control and induction values, respectively) and 24 h after the first and fourth RT-fractions in weeks 1, 2, 4 and 6 (after fractions 1, 4, 6, 9, 16, 19, and 26; and after fraction 29 in head and neck cancer samples).

To perform TEM-analysis, blood samples were collected directly before and 0.1, 0.5 and 24 h after the first RT-fraction for immediate processing.

For *ex-vivo*-experiments, blood from healthy donors was obtained, PBLs isolated, homogeneously irradiated, and incubated in RPMI at 37°C.

Dose dependence: PBLs were suspended in cold phosphate-buffer saline (PBS), irradiated with different doses (0.5, 1.0, 2.0 or 4.0 Gy), and suspended in RMPI-Medium (Sigma-Aldrich, St. Louis, MO) prior to a 0.5 h incubation at 37°C allowing for repair.

Time course: Following irradiating with 1.0 Gy, PBLs were incubated in RPMI-Medium at 37°C, and fixated 0.1, 0.25, 2.5, 8.0 and 24 h after irradiation. Non-irradiated PBLs from the same donor served as control.

### Blood sample preparation for IFM and TEM

Briefly, blood samples in heparin tubes were diluted with 6 ml RPMI and incubated at 37°C. PBLs were isolated using a kit (PAA Laboratories; Cölbe, Germany). Blood samples were layered on Percol 400 and centrifuged at 1200 g for 20 minutes. 5 ml PBS was added to the resulting interphase and centrifuged at 300 g for 10 minutes. The separation yielded ~80% PBLs, ~15% monocytes and ~5% granulocytes.

For IFM, PBLs were fixed in 100% methanol for 0.5 h and permeabilized in 100% acetone for 1 minute at −20°C. After washing cells in PBS with 1% fetal calf serum for 1 x 10 minutes at room temperature, samples were incubated with 53BP1 antibody (anti-53BP1, mouse monoclonal, Merck, Darmstadt, Germany) followed by a secondary fluorescent antibody (AlexaFluor-488, Invitrogen, Karlsruhe, Germany). Samples were mounted using Vectashield mounting medium (Vector Laboratories, Burlingame, CA) with DAPI (4’,6-Diamidino-2-phenylindole). Fluorescent images were captured and visually analyzed. A trained staff member identified and counted the cells until at least 300 cells and 40 foci for each time point were registered. All PBLs in each field of view were analyzed, even those without evidence of radiation damage.

For TEM, PBL pellets were fixed overnight with 2% paraformaldehyde and 0.05% glutaraldehyde in PBS. The ethanol-dehydrated samples were infiltrated with LR Gold resin (EMS, Hatfield, PA). Afterwards, samples were embedded in resin containing 0.1% benzyl and kept for 24 h at −20°C followed by ultraviolet light exposure until resin was polymerized. Ultrathin 70 nm slices were sectioned off the samples using a Microtome Ultracut UCT (Leica, Biel, Switzerland), picked up on pioloform-coated nickel grids, and processed for immunogold labeling. To block non-specific staining, sections were floated on drops of 50 mM glycine and blocking solution. Afterwards, following rinsing, sections were incubated with different primary antibodies (53BP1 or pKu70 (anti-pKu70, rabbit polyclonal, pSer5, Abcam, Cambridge)) overnight at 4°C. The same primary antibodies used in fluorescence microscopy were applied in combination with gold-labeled secondary antibodies for TEM experiments in order to visualize pKu70 and detect incomplete DNA damage repair sites. A single IFM-focus has a diameter of approximately 1.0 μm (S1A Fig; green circle). When using gold-labeled secondary antibodies in the same IFM-approach, this focus consists of two 53BP1-clusters, each one with a diameter of only 500 nm (S1B Fig; red circles). In TEM analysis, this focus-area can be subdivided further into euchromatic and heterochromatic compartments (S1C Fig) and thus allows for reliable detection and quantification of DNA repair factors (S1D Fig; pKu70, 10 nm, gold beads colored in red; 53BP1, 6 nm, colored in green) and their localization within different chromatin compartments.

After rinsing, goat secondary antibodies conjugated with 6 nm and 10 nm gold particles (EMS) were applied to the sections on the grids, and then incubated for 1.5 h at room temperature. Subsequently, sections were washed and fixated with 2% glutaraldehyde in PBS. All sections were stained with uranyl acetate and examined with a Tecnai Biotwin™ transmission electron microscope (FEI, Eindhoven, Netherlands). For quantification, we identified pKu70-dimers (two 10 nm gold particles) and 53BP1 bead-clusters (6 nm) visually at 48000-86000x magnification and counted these in 50 randomly chosen nuclear sections.

### Statistical analysis

A one sided Mann-Whitney Test was performed using the statistical software OriginPro (version 8.5, OriginLab Corporation, Northampton, USA) to evaluate potential differences between data groups. The criterion for statistical significance was p ≤ 0.05.

The dispersion index test was used to determine the deviation of foci per cell-distribution at the 0.5 h data point from Poisson statistics to demonstrate that – in the setting of partial body irradiation to the head and neck or pelvic region – only a proportion of PBLs was exposed to irradiation [28, 29]. The test was performed with the software Dose Estimate, version 3.0 (Chilton, United Kingdom).

## Results

The study population consisted of nine individuals with head and neck or rectal cancers (5 patients received RCT and 4 RT without chemotherapy). Table 1 shows the patients’ characteristics. Based on our assumption that the number of irradiation induced DSBs depends on the applied dose and irradiation time, patients were grouped according to their cancer type and the technique applied (patients with head and neck-cancers received IMRT while those with rectal-cancers underwent 3D-CRT; S2A and S2B Figs). To show the influence of chemotherapy on DSB-formation, patients were further divided into those who received chemotherapy and those who did not. In total, 40 samples from patients with head and neck cancer and 46 from patients with rectal cancer (three technical replicates per sample) were analyzed.

**Table 1.** Patients and treatment-characteristics according to cancer type.

| Category | head&neck cancer | rectal cancer |
| --- | --- | --- |
| **Total no. of patients** | 4 | 5 |
| **Age, years** |  |  |
| mean ± SD | 64 ± 5 | 60 ± 10 |
| range | 60 - 71 | 50 - 74 |
| **Sex, no. (%)** |  |  |
| Male | 3 (75) | 3 (60) |
| Female | 1 (25) | 2 (40) |
| **KPS, no. (%)** |  |  |
| 70 |  | 1 (20) |
| 80 | 4 (100) | 2 (40) |
| 90 |  | 1 (20) |
| 100 |  | 1 (20) |
| **T-Stage, no. (%)** |  |  |
| T1 | 1 (25) |  |
| T2 | 3 (75) |  |
| T3 |  | 4 (80) |
| T4 |  | 1 (20) |
| **N-Stage, no. (%)** |  |  |
| N0 | 2 (50) | 2 (40) |
| N1 |  | 1 (20) |
| N2 | 2 (50) | 2 (40) |
| **Chemotherapy\*, no. (%)** |  |  |
|  | 1 (25) | 4 (80) |
| **Irradiation technique** |  |  |
|  | IMRT<br>(6 MV photons) | 3D-CRT<br>(18 MV photons ) |
| **Irradiation time per fraction** |  |  |
| mean | 30 min | 7 min |
| **Total dose, no (%)** |  |  |
| 46.8 Gy |  | 1 (20) |
| 50.4 Gy |  | 4 (80) |
| 60.0 Gy | 4 (100) |  |
| **PTV, cm<sup>3</sup>** |  |  |
| mean ± SD | 1047 ± 226 | 1411 ± 444 |
| **D90-PTV, % ref.-dose ± SD** | 89 ± 6 | 94 ± 5 |
| **D80-PTV, % ref.-dose ± SD** | 95 ± 2 | 97 ± 3 |
| **Isodose volume, cm<sup>3</sup>** |  |  |
| V5<sub>iso</sub> ± SD | 6703 ± 1876 | 11961 ± 1411 |
| V10<sub>iso</sub> ± SD | 5184 ± 1447 | 10030 ± 1012 |
| V20<sub>iso</sub> ± SD | 3872 ± 924 | 7099 ± 647 |
| V30<sub>iso</sub> ± SD | 2704 ± 627 | 5575 ± 510 |
| V45<sub>iso</sub> ± SD | 1427 ± 384 | 2183 ± 703 |
| **Blood volume<sup>#</sup>, cm<sup>3</sup> ± SD** |  |  |
| | 117 ± 27 | 173 ± 44 |

SD: standard deviation. KPS: Karnofsky Performance Score. PTV: Planning Target Volume.
\*Concurrent chemotherapy regime as described in 'Patients and Methods', three patients with rectal cancer received neoadjuvant regimen, one patient received adjuvant regimen.
<sup>#</sup>Derived from delineated blood vessels >1cm in diameter and encompassed by the 50%-isodose line.

Quantification of initial foci induction by IFM was completed on samples taken 0.5 h after the first RT-fraction. 53BP1-foci were not detected in 39 of 432 PBLs (~10%) from patients with head and neck cancer and in 50 of 517 (~10%) from those with rectal cancer, confirming that partial body irradiation causes limited PBL exposure (Table 2). In contrast, no 53BP1-foci could be detected in 85% of the unirradiated PBLs (in total 2735 from 3222 cells) taken before the first fraction.

**Table 2.** Dispersion analysis of 53BP1-foci distribution 0.5 h after the first RT-fraction as measured by immunofluorescence.

| Tumor entity | 53BP1-yield, foci per cell |  |  |  |  |  |  |  |  |
| --- | --- | --- | --- | --- | --- | --- | --- | --- | --- |
|  | 0 | 1 | 2 | 3 | 4 | 5 | 6 | 7 | ≥8 |
| **Head&neck cancer, no. of cells** |  |  |  |  |  |  |  |  |  |
| Observed distribution* | 39 | 60 | 61 | 91 | 55 | 52 | 26 | 21 | 27 |
| Poisson distribution | 14 | 48 | 82 | 94 | 80 | 56 | 32 | 16 | 4 |
| **Rectal cancer, no. of cells** |  |  |  |  |  |  |  |  |  |
| Observed distribution# | 50 | 66 | 103 | 100 | 81 | 47 | 32 | 17 | 21 |
| Poisson distribution | 22 | 68 | 108 | 114 | 92 | 58 | 32 | 14 | 8 |

\* 432 PBLs were analyzed in four patients. The resulting distribution significantly deviates from a Poisson distribution, indicating a partial body irradiation [mean dispersion index is $1.8 \pm 0.1$ (standard error of the mean), U value (standard normal deviate) is 8.3, and irradiated fraction of cells is 93% as calculated with the contaminated Poisson method].

To compare the appearance of 53BP1-foci among samples from different treatment types, we looked at the PTV-size, radiation duration and exposed blood volume from delineated blood vessels (>1 cm diameters) encompassed by the 50%-isodose line. Table 2 shows the 53BP1-foci distribution analysis results measured by IFM 0.5 h after the first RT-fraction.

The 53BP1-foci distribution of homogeneous *ex-vivo*-irradiated PBLs follows the Poisson statistic [30, 31] as shown in Table 2. A linear dose response relationship was found 0.5 h after irradiation (0.5 - 4.0 Gy) (Figs 1A and 1C) and in 53BP1-foci loss 24 h post exposure (1.0 Gy) (Figs 1B and 1D).

**Fig 1.**
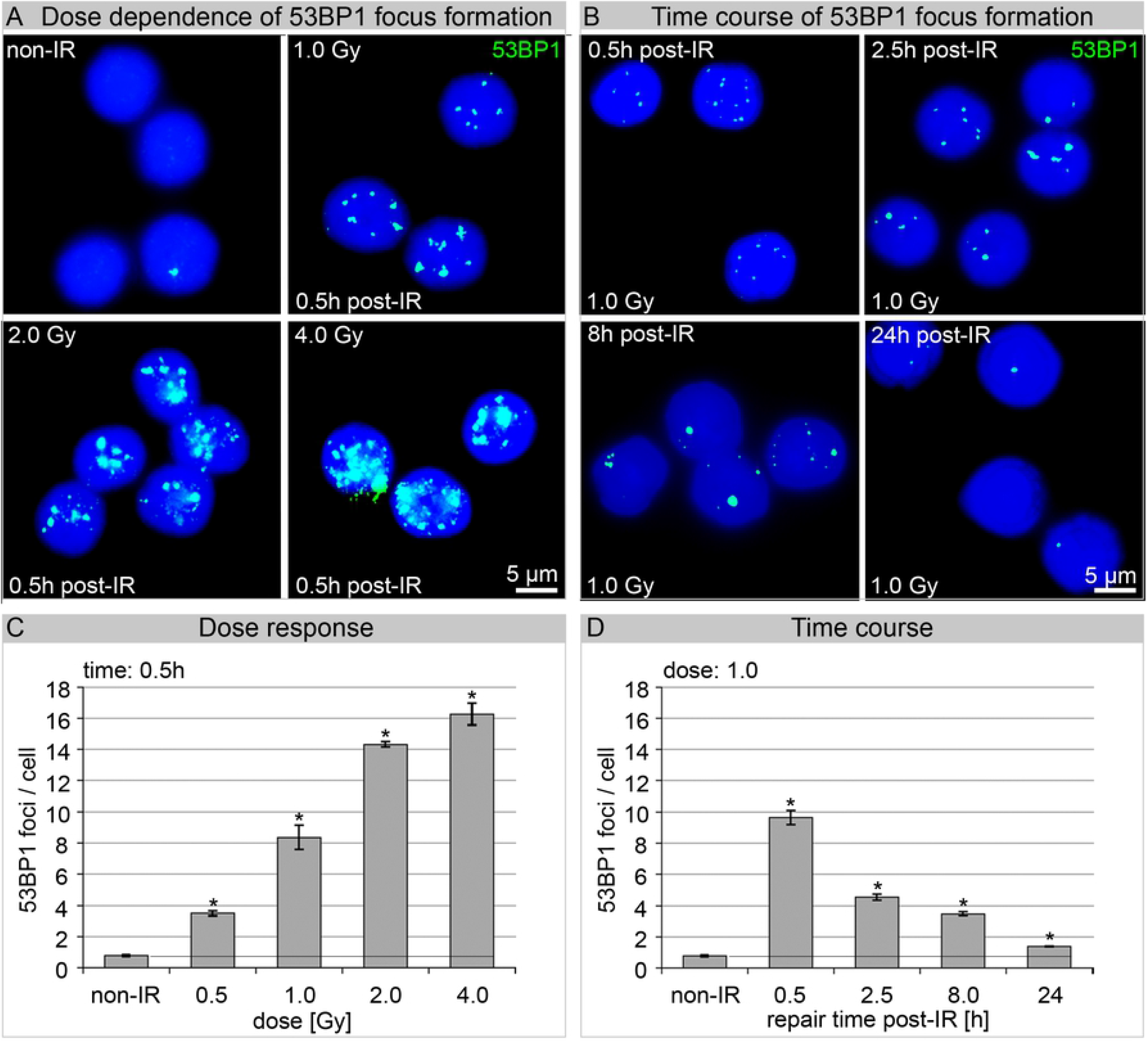
Dose dependence and time course of 53BP1 focus formation. (A) Immunofluorescence-staining for 53BP1 in PBLs analyzed before (non-IR) and 0.5 h after homogeneous irradiation with 1.0, 2.0 or 4.0 Gy. (B) Time kinetics of radiation-induced 53BP1-foci. (C,D) 53BP1 were visually counted as number of foci/cell. All points are mean values of three different experiments where at least 300 cells were counted from 10 randomly chosen fields of view [* statistically significant differences (p ≤0.05) compared with non-irradiated controls and previous values].

Fig 2A shows the amount of 53BP1-foci per cell counted at each time point for all patients stratified by tumor entity using IFM.

**Fig 2.**
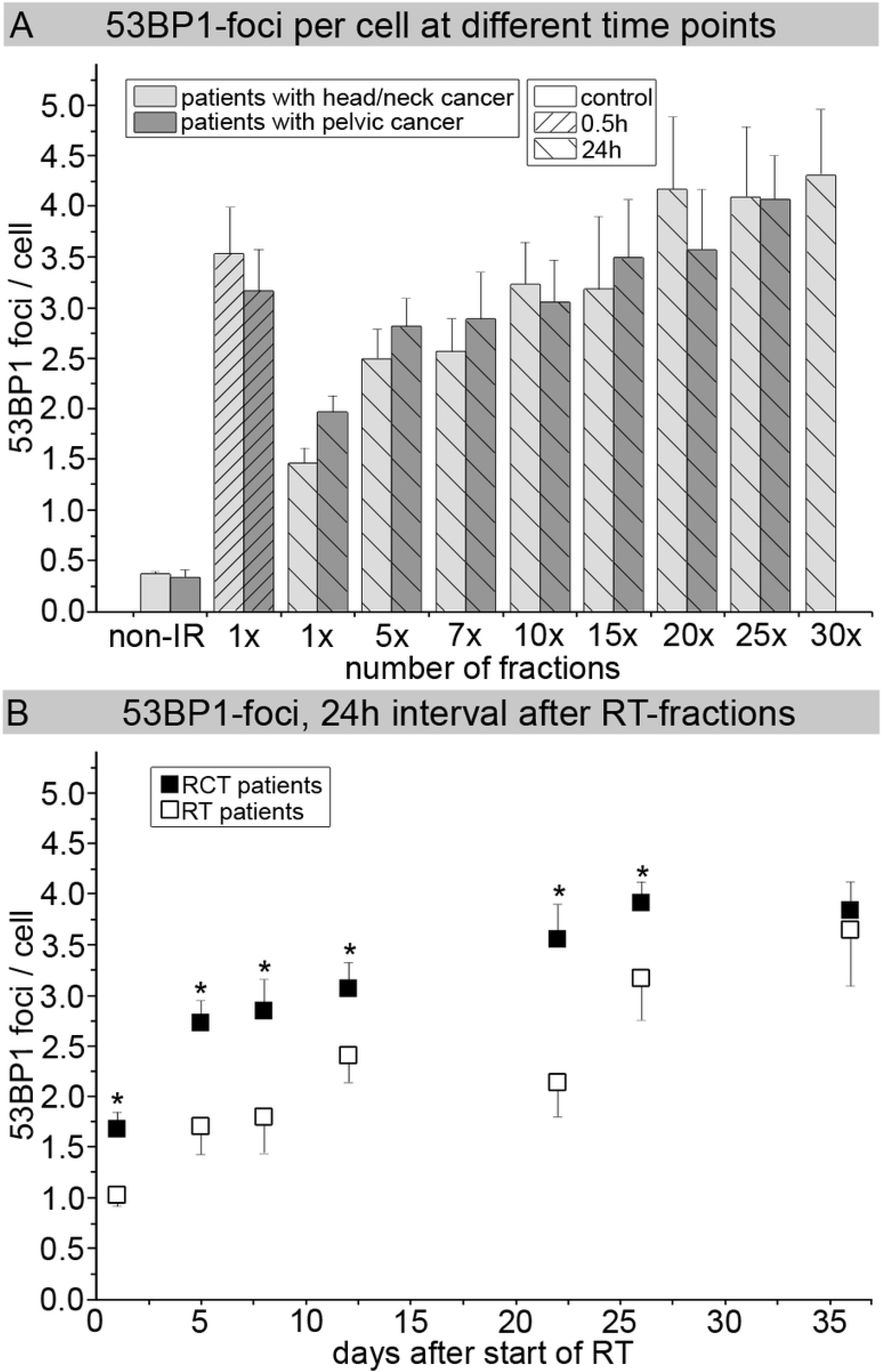
Quantification of 53BP1-foci by IFM. (A) The number of 53BP1-foci per PBL nucleus was counted before (non-IR), 0.5 h and 24 h after the first dose fraction (1x) as well as, 24 h after a predefined number of additional *in-vivo* irradiation fractions (5x to 30x) to the head and neck (n = 4) or rectal (n = 5) regions. Data are presented as mean values of three technical replicates per patient ± standard error. (B) 24h-data during fractionated RT progression, stratified by the administration of concurrent chemotherapy. Data are presented as mean values of three technical replicates per patient ± standard error; the background number of foci/cell (control in week 1 before irradiation) was substracted from the data. * Significant difference to RT patients (p ≤0.05).

The control showed a low number of foci in patients with head and neck cancer (0.37 ± 0.02 53BP1-foci/cell) and in those with rectal cancer (0.33 ± 0.08 53BP1-foci/cell). At 0.5 h after the first RT-fraction, we found a 10-fold rise in the 53BP1-foci number in head and neck cancer patient samples (3.54 ± 0.46 53BP1-foci/cell) and rectal cancer patient samples (3.17 ± 0.40 53BP1-foci/cell). Over time, PBLs from both groups showed a decrease in foci numbers, although it was still possible to visualize an average number of 1.46 ± 0.05 53BP1-foci/cell in PBLs from head and neck cancer patients and 1.97 ± 0.17 (~62%) 53BP1-foci/cell in PBLs from rectal-cancer patients 24 h after the first fraction (1x) (Fig 2B).

With increasing numbers of administered RT-fractions (up to 30x), the 53BP1-foci accumulated. Patients with head and neck cancer had 2.49 ± 0.30 53BP1-foci/cell (5x) 24 h after the first week of RT-fractions and 4.31 ± 0.65 53BP1-foci/cell (30x) 24 h after further irradiation. Patients with rectal-cancer had 2.81 ± 0.29 53BP1-foci/cell (5x) and 4.07 ± 0.44 53BP1-foci/cell (25x), respectively (Fig 2A).

The 53BP1-foci-levels in PBLs of patients with head and neck cancer tended to outnumber those of patients with rectal cancer; however, the differences were not significant. Moreover, we analyzed the number of 53BP1-foci according to whether patients received accompanying chemotherapy (Fig 2B). Patients receiving chemotherapy had 1.69 ± 0.16 53BP1-foci/cell, 24 h after the first RT-fraction (1x) and those without chemotherapy had 1.02 ± 0.04 53BP1-foci/cell at the same time point. During the course of therapy, 53BP1 accumulated to 2.74 ± 0.08 foci/cell (5 days after start of RT) up to 3.78 ± 0.34 foci/cell (36 days after start of RT). Without accompanying chemotherapy, 53BP1-foci values were significantly lower with 1.70 ± 0.24 53BP1-foci/cell (5 days after start of RT) up to 3.17 ± 0.86 53BP1-foci/cell (26 days after start of RT).

The quantification of 53BP1-foci through IFM enables the estimation of DNA-repair capacity after irradiation exposure. Application of TEM-analysis improved the resolution of DNA damage patterns which are obscured in IFM by the fluorescence of the labeled foci. Quantification of pKu70 and 53BP1 in PBLs after homogeneous *ex-vivo* irradiation verified the suitability and reliability of the TEM-method. Immunogold labeling of PBLs for pKu70 (10 nm bead-size, colored in red) and 53BP1 (6 nm, colored in green), was completed 0.5 h after 1.0 Gy irradiation. Colocalization of 53BP1-clusters with pKu70-dimers was observed exclusively in heterochromatic areas. Additionally, pKu70-single-beads and small 53BP1-clusters (2 to 5 beads) were occasionally present at the border between euchromatic and heterochromatic domains (Fig 3A).

**Fig 3.**
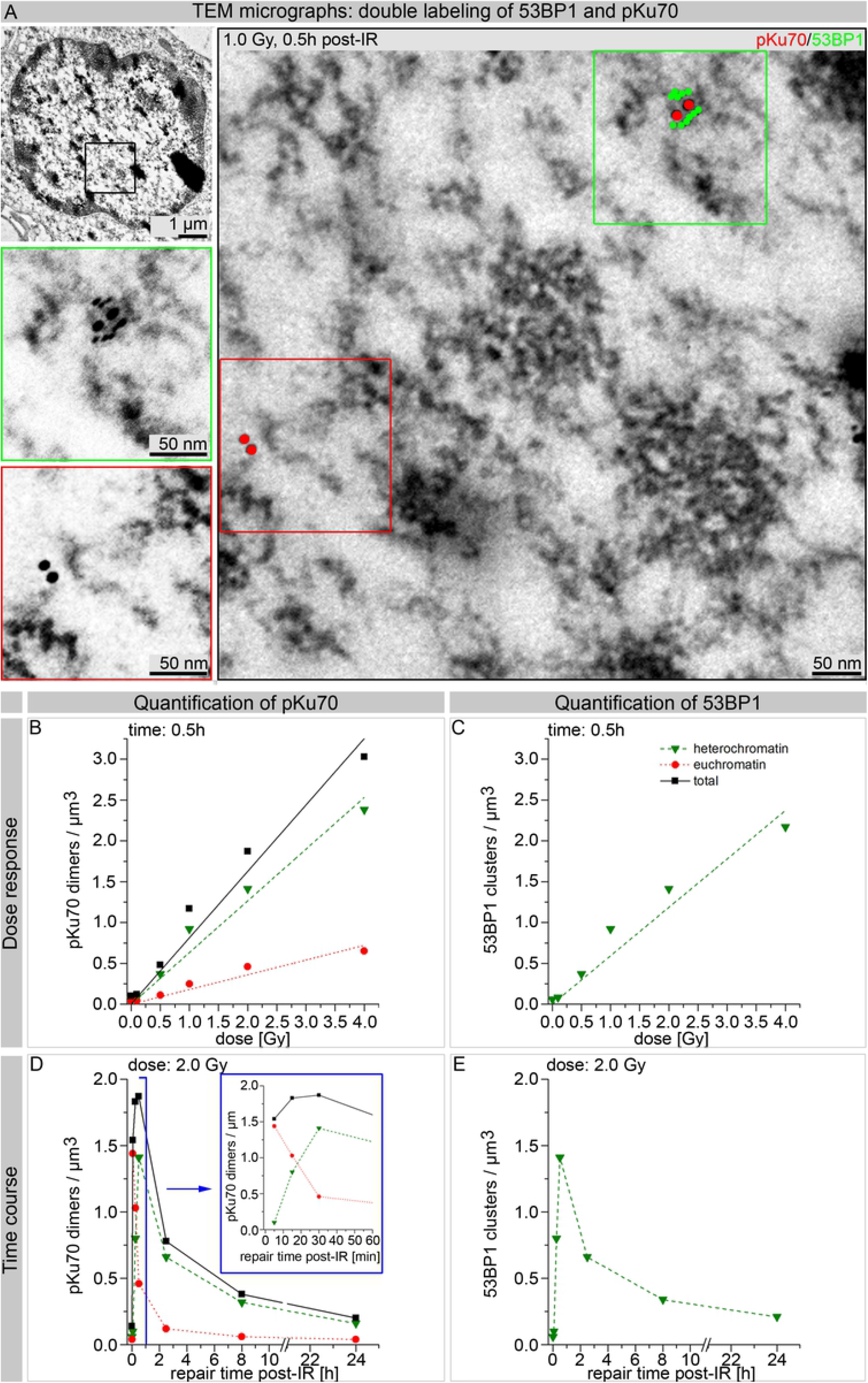
Quantification of pKu70-dimers and 53BP1-clusters by TEM after homogeneous irradiation with 1.0 Gy. (A) Visualization of pKu70 (red, 10 nm beads) and 53BP1 (green, 6 nm) 0.5 h after 1 Gy irradiation in a representative TEM-image. Framed regions shown at higher magnification in adjacent images. Quantification of pKu70-dimers and 53BP1-clusters (quantified in ≥ 50 nuclear sections) in eu- and heterochromatin 0.5 h after irradiation with different doses (B, C) and at different time points after irradiation with 2 Gy (D; E).

In line with previous data [32], quantification of a dose response (0.1, 0.5, 1.0, 2.0 or 4.0 Gy) 0.5 h after irradiation revealed that the total number of pKu70-dimers and 53BP1-clusters was dose-dependent (Figs 3B and 3C). Gold bead dimers and clusters were normalized to nuclear area and section thickness (pKu70-dimers/μm^3^ or 53BP1-clusters/μm^3^).

A straight-line correlation from 0.12 pKu70/μm^3^ (1.0 Gy) up to 3.03 pKu70/μm^3^ (4.0 Gy) with a background signal of 0.10 pKu70/μm^3^ in non-irradiated PBLs was demonstrated. The number of induced pKu70-dimers in euchromatin was consistently lower (from 0.04 pKu70/μm^3^ at 0.1 Gy to 0.65 pKu70/μm^3^ at 4.0 Gy) than those reached in heterochromatin (from 0.08 pKu70/μm^3^ to 2.38 pKu70/μm^3^). This indicates that 0.5 h after irradiation, portions of the originally induced euchromatic DNA damage was no longer present, whereas, the highest recognition of heterochromatic DSBs took place at this time point. The number of 53BP1-clusters and pKu70-dimers almost correlate completely (from 0.08 53BP1/μm^3^ to 2.17 53BP1/μm^3^) due to their colocalization.

Analysis of the time kinetics occurred in the same manner, only at different time points (0.1, 0.25, 0.5, 2.5, 8.0 and 24 h) after homogeneous irradiation with 1.0 Gy. Highest values of pKu70-dimers were observed in euchromatic compartments after 0.1 h (1.44 pKu70/μm^3^). Ultimately, this value decreased to 1.03 pKu70/μm^3^ (~72%) after 0.25 h, to 0.46 pKu70/μm^3^ (~32%) after 0.5 h, to 0.12 pKu70/μm^3^ (~8%) after 2.5 h, to 0.06 pKu70/μm^3^ (~4%) after 5 h and to 0.04 pKu70/μm^3^ (~3%) after 24 h (Fig 3D). These results suggest that euchromatic DSBs are quickly recognized following irradiation and can be completely repaired within a few hours. On the contrary, the number of heterochromatic pKu70-dimers initially rose from 0.10 pKu70/μm^3^ (0.1 h post-IR) to 0.80 pKu70/μm^3^ (0.25 h post-IR) and to 1.41 pKu70/μm^3^ (0.5 h post-IR). Subsequently, the pKu70-dimers began to decrease in numbers 2.5 h after irradiation to 0.66 pKu70/μm^3^ (~47%), to 0.32 pKu70/μm^3^ (~23%) after 8 h, and to 0.16 pKu70/μm3 (~11%) after 24 h. The 53BP1-clusters showed roughly the same kinetics as the heterochromatic pKu70-dimers with an increase from 0.10 53BP1/μm^3^ (0.1h post-RT) to 0.80 53BP1/μm^3^ (0.25h post-IR), and 1.41 53BP1/μm^3^ (0.5h post-IR). Then, 0.66 53BP1/μm^3^ (~47%) 53BP1-clusters were detectable after 2.5 h and decreased to 0.34 53BP1/μm^3^ (~24%) after 8 h, and 0.21 53BP1/μm^3^ (~15%) were still visible after 24 h (Fig 3E). Additionally, 53BP1-clusters, consisting of 5 to 12 gold beads and without pKu70 colocalization, were observed 8 h and 24 h after irradiation, potentially marking chromatin changes in areas where DNA-damage was present.

To expand our knowledge on DNA-damage, we investigated PBLs of patients with head and neck cancer after RT and RCT using TEM, before and 0.1, 0.5 and 24 h after the first fraction by quantifying pKu70-dimers and 53BP1-clusters in euchromatin and heterochromatin of 50 nuclear sections per sample. Figs 4A and 4B show representative TEM-micrographs. Visualization of 10-nm (pKu70-dimers) and 6-nm-gold beads (53BP1-cluster) was improved by overlaying with red and green circles respectively. The RCT-patient (Figs 4C-F) showed a higher level of repair proteins than the RT-patient in both chromatin domains (Fig 4C) 0.5 h after the first fraction.

**Fig 4.**
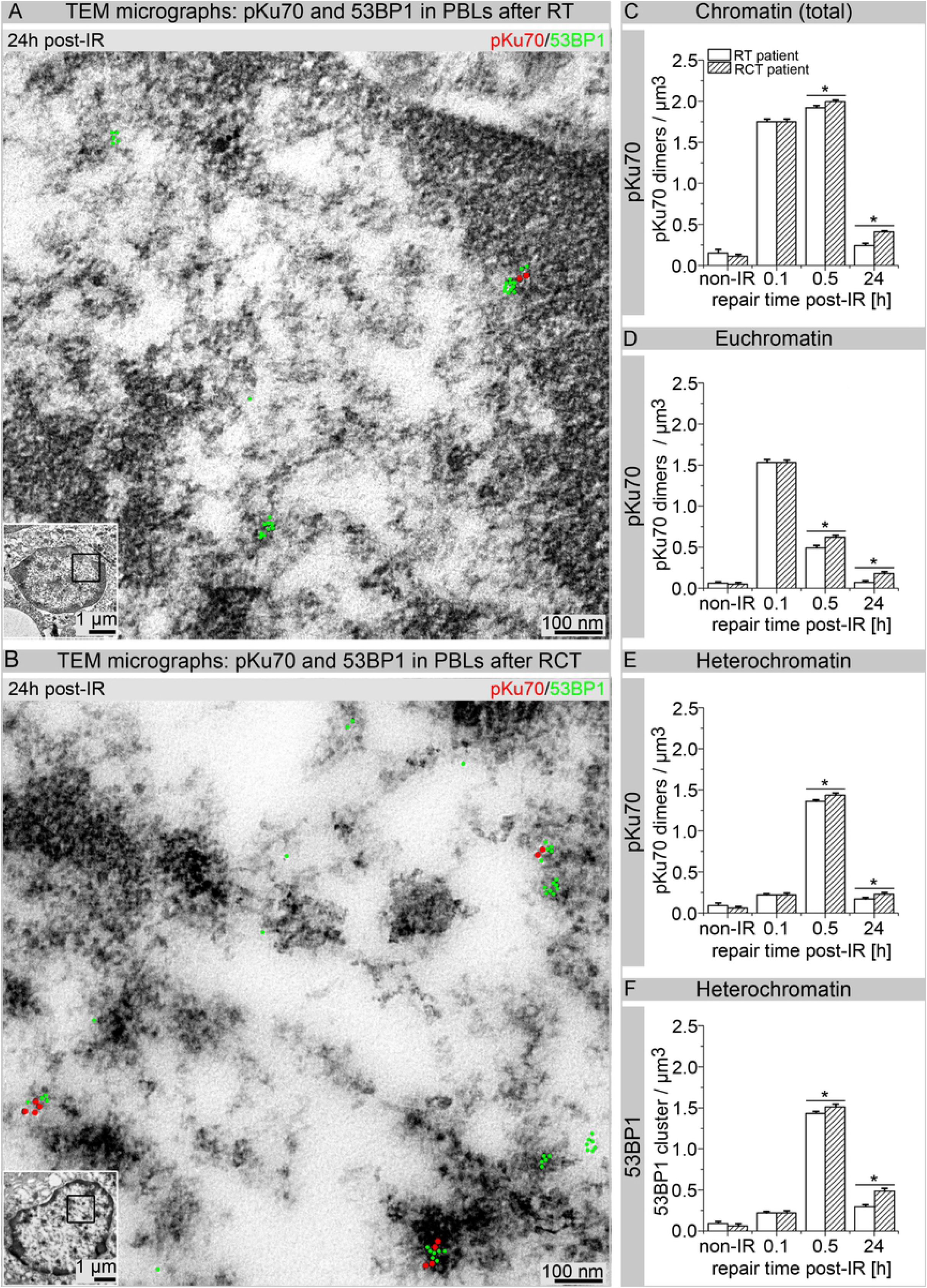
Quantification of pKu70-dimers and 53BP1-clusters by TEM in PBLs of patients who received RT or RCT. Visualization of pKu70 and 53BP1 in peripheral blood lymphocytes (PBLs) 24 h after first RT (A) and RCT (B). TEM quantification of pKu70-dimers per μm^3^ in total chromatin (C), euchromatin (D) and heterochromatin (E) following RT or RCT. Additionally, the number of 53BP1-beads per μm^3^ was quantified in PBLs of RT and RCT patients 0.1, 0.5, and 24 h after the first radiation fraction.

Controls showed a low level of pKu70-dimers/μm^3^ (RT: 0.15 ± 0.01 pKu70-dimers/μm^3^; RCT: 0.11 ± 0.01 pKu70-dimers/μm^3^). 0.1 h after irradiation no significant differences were found in pKu70 between RT and RCT (RT: 1.75 ± 0.03 pKu70-dimers/μm^3^; RCT: 1.76 ± 0.04 pKu70-dimers/μm^3^). A similar pattern for 53BP1-clusters could be seen (Fig 4F), with significant higher values for RCT-patients at 0.5 h and 24 h after the first irradiation (S1 Table).

## Discussion

In this study, we questioned whether the DNA damage repair in PBLs during radiotherapy for head and neck or rectal cancers is influenced by simultaneous chemotherapy or other variables, such as isodose volume, irradiation time, or by different irradiation techniques (IMRT or 3D-CRT). Additionally, we investigated how repetitive heterogeneous dose exposure influences the radiation-induced DNA damage of PBLs, especially in patients prescribed with concomitant chemotherapeutics. To do this, we quantified 53BP1-foci-formation [8, 9] in *ex-vivo*-irradiated PBLs, and found that not every 53BP1-focus equates to an unrepaired DSB.

IFM is a well-established method to visualize and analyze DNA repair proteins. It has the advantage of allowing fast examination of many cells. However, this is only possible for repair factors that accumulate in the vicinity of DSBs in sufficient amounts thereby producing adequate fluorescent signals. IFM cannot be used to detect pKu70, which binds as a heterodimer with pKu80 at the ends of DSBs [4, 32]. Visualization of 53BP1-foci using IFM indicates DSBs, however persisting 53BP1-foci mean fluorescence signals may still be detectable after the initial damage has been repaired. TEM enables clarification of the repair status of a DSB as it allows for pKu70 detection, the central repair protein of the NHEJ, which signals incomplete DNA damage repair.

Several factors or modified proteins can be employed to analyze DSBs. One commonly used factor is the phosphorylated histone protein 2A (γH2AX), and another is 53BP1. We chose 53BP1 as after DSB induction, γH2AX is found along several megabase pairs (Mbp) of the DSB [33], whereas 53BP1 gets recruited to the immediate vicinity of a DSB [34–37]. While the larger γH2AX areas are advantageous for IFM as they can be easily detected, they are not the best choice when applying TEM due to its higher resolution and the need for combined immunogold labeling. As a result, TEM displays γH2AX molecules as linear gold bead accumulations (called bead chains) [24]. Due to the three-dimensional nature of the nucleus, these bead chains are not found as parallel chains on the surface of the investigated section. Hence, only parts of the chain can be visualized in each individual section. As a consequence, allocation of γH2AX chain-sections to a single DSB (represented by a pKu70-dimer) in TEM is difficult, especially when several pKu70-dimers are present. On the contrary, 53BP1 is better suited for TEM since it is detectable locally at the DSB shortly after irradiation and independently as persisting 53BP1-clusters without pKu70-colocalization (Figs 4A and 4B).

Following homogeneous PBL irradiation, IFM showed a linear 53BP1 dose correlation up to 1 Gy (Fig 1C) and TEM showed similar correlation up to 4 Gy (Fig 3C). With increasing doses (> 1 Gy), the fluorescent signal of individual adjacent foci overlapped, making quantification in IFM difficult or almost impossible (Fig 1A). This limitation does not occur with quantitative TEM as gold beads were either present and quantifiable or absent. However, when LR-gold-resin embedded PBL sections are investigated and quantified using TEM it is critical to always consider that only plane sections of the nucleus and not the entire cell nucleus are examined. To gain greater insight in the number of the gold-labeled repair proteins in the entire nucleus, labeling in 50 nuclei sections per dose or repair time point was quantified. Additionally, non-homogeneously distributed pKu70-dimers and 53BP1-clusters in the cell nucleus over the number of nuclei sections was captured as reliably as possible by counting all 10-nm (pKu70) and 6-nm (53BP1) gold beads. The distance of a 10-nm gold bead from the antigen or repair protein is a maximum of 28 nm (S3 Fig). In contrast, IFM enabled visualization and quantification of foci in each entire PBL nucleus. The primary antibodies and the fluorochrome-coupled secondary antibodies penetrated the fixed cells and cell nuclei due to the permeabilization step (acetone, 1 min at −20 ° C). In IFM, the degree to which cell structures were preserved after this chemical treatment was not detectable by means of the DAPI-signal. No other publications have reported on the structural preservation and quality of cells after IFM sample preparation. After sectioning embedded PBLs immunogold-labeled repair proteins were visualized and quantified in TEM. Permeabilization was not necessary, as antigens were found freely accessible near the surface. All cell structures (membranes, mitochondria, etc.) were perfectly visible in TEM and optimally preserved.

PBLs, which are often used for biological dosimetry [38] and for the determination of individual radio-sensitivity [39], do not go through the cell cycle but remain in the G0-phase. Several studies have shown that PBLs sometimes undergo apoptosis 12-24 h after irradiation, which is characterized by significant chromatin condensation [40, 41]. This apoptotic chromatin condensation within human PBLs may prevent the decomposition of residual DNA-repair-foci, which were observed in PBLs 24 h after irradiation [42]. Irradiation-induced residual foci in condensed chromatin can persist longer than 24 h in apoptotic G0-PBLs. Persisting foci are not to be confused with existing DSBs as these especially slow repaired or unrepaired DSBs are eventually responsible for induction of apoptosis. Moreover, pulsed-field gel electrophoresis and confocal laser microscopy experiments have shown that in normal human fibroblasts, repair of existing DSBs does not correspond with the counted 53BP1-foci [43]. Our results confirm this, as 24 h after irradiation 53BP1-clusters often did not colocalize with pKu70 (Figs 4A and 4B). As we have already reported, DSBs can be visualized by pKu70-dimers [25, 32] that bind directly to the ends of DSBs. In TEM, lack of pKu70 within a 53BP1-cluster indicates the absence of a DSB at this point. Therefore, the frequency of colocalizations between pKu70 and 53BP1 is largely dependent on the point in time post-irradiation. The recorded accumulation of 53BP1-foci by IFM, in Fig 2A, after an increasing number of applied fractions, is probably due to a mixture of newly induced DSBs per daily fraction and the generation of persistent chromatin changes after unfinished or defective DSB-repairs. Loss of 53BP1-foci occurs through the elimination of aged or damaged PBLs and the successful repair of DSBs. However, as we observed an overall increase in the number of 53BP1-foci, especially 24 h post-IR between the first (1x) and last (25x or 30x) fraction, it is reasonable to assume that at this time point accumulation of persisting 53BP1-foci was significantly involved, whereas during the previous 24 h more repairable DSBs were eliminated. Thus, it is not possible to evaluate the repair capacity based on a rise in 53BP1- foci detected 24 h after irradiation. Most persistent 53BP1-foci responsible for the increase are located exclusively in the periphery of heterochromatin domains and contain no pKu70. Most likely, these represent apoptotic processes rather than unrepaired DSBs. In Fig 2B, we show higher 53BP1-foci-levels for all patients who received RCT, independent of the collective.

By using the higher resolution of TEM in combination with the immunogold-labeling of pKu70 and 53BP1 within the intact nuclear cell ultrastructure, we were able to detect a higher number of pKu70-dimers in the euchromatin and in the heterochromatin of PBLs in RCT-patients than in those of patients after a single RT 24 h post-IR (Fig 4). In addition, we visualized pKu70-dimers individually as well as collectively (2x or 3x pKu70-dimers) which suggests multiple DSBs in close proximity (Fig 4B). These results indicate that the scale of DNA-damage induced during radiotherapy is affected by the presence of accompanying chemotherapy. Thus, the number of DSBs in PBLs was not significantly influenced by the irradiation technique (IMRT or 3D-CRT) or the size of the irradiation field. This, however, could be due to the possibility that effects induced by IMRT, in which a smaller PTV is irradiated over a longer period of time, counterbalance those induced in 3D-CRT in which a larger volume is irradiated over a shorter time-period.

Based on these data, we propose that persisting DSBs (pKu70-dimers) represent more severe damage induced by RCT (1, 2 or more pKu70-dimers representing multiple DNA-lesions). Repair seems to be difficult or even impossible in these cells. In addition, as persistent DSBs were detectable only at certain times >0.5 h after irradiation and always at the edge of heterochromatic domains, we suspect that cellular processes, such as the opening of densely packed heterochromatic regions containing one or more DSBs, delay repair. However, not every residual focus is equivalent to a remaining DSB, since pKu70 was not present at every damage site. Persistent 53BP1-clusters without colocalizing pKu70 are likely to show chromatin alterations after completion or possibly, defective repair. Therefore, IFM-53BP1-foci-analyses alone are not adequate to determine the individual repair capacity after the irradiation of PBLs, as a DSB may be marked by a 53BP1 focus but not every 53BP1 focus represents a DSB.

## Acknowledgments

The authors thank Liz Ainsbury and Kai Rothkamm for providing the software Dose Estimate 3.0 and helpful discussions, Patrick Melchior, M.D., for his valuable support in the statistical analysis of foci yield curves, Sara Timm and Nadine Schuler for excellent technical assistance in IFM analysis, and Anna Isermann for editorial assistance in the preparation of the manuscript.

## Data Availability

All relevant data are within the paper and its supporting information files.

## Author Contributions

CER and YL conceived and designed the experiments. YL and PG performed experiments. CER, YL and JF analyzed the data. CER and CR contributed with reagents, materials and analysis tools. YL wrote the paper. CER and JF reviewed the manuscript.

## Supporting information

**S1 Fig.** Different resolution powers of light- and electron microscopy.

(A) Immunofluorescent image of 53BP1-foci, 0.5 h after irradiation with 1 Gy in the DAPI-stained nucleus of a peripheral blood lymphocyte. pKu70 cannot be observed by IFM.

(B) Light microscopy image of a PBL-nucleus. By virtue of antibodies targeting 53BP1, two clusters can be seen (red circles).

(C) Electron microscopy image (TEM, 2700x magnification). Euchromatin (bright) and heterochromatin (dark) can be clearly differentiated within the nucleus.

(D) Reliable visualization of immunogold-labeled 53BP1 (green) and pKu70 (red) by means of TEM (48 000x).

**S2 Fig.** Representative isodose distribution. (A) IMRT was mandatory for patients with head and neck cancer with a pre-defined arrangement of seven coplanar beam angles. (B) For patients with rectal cancer through 3D-CRT with three coplanar portals. The irradiated volume within the violet isodose (10% reference dose), varies in size, depending on the cancer type and irradiation modes.

**S1 Table.** PBLs of patients with head and neck cancer after RT and RCT were investigated using TEM, before and 0.1, 0.5 and 24 h after the first fraction by quantifying pKu70-dimers and 53BP1-clusters in euchromatin and heterochromatin of 50 randomly chosen nuclear sections. Gold bead dimers and clusters were normalized to nuclear area and section thickness (pKu70-dimers/μm^3^ or 53BP1-clusters/μm^3^) and presented as mean values.

**S3 Fig.** 3D-model (using *computer-aided design*, AutoCAD 2017, Autodesk GmbH, USA) of a primary antibody, bound to a secondary antibody coupled to a 10 nm colloidal gold particle.

